# Highly efficient transgenic mouse production using piggyBac and its application to rapid phenotyping at the founder generation

**DOI:** 10.1101/2023.12.10.570953

**Authors:** Eiichi Okamura, Seiya Mizuno, Shoma Matsumoto, Kazuya Murata, Yoko Tanimoto, Dinh Thi Huong Tra, Hayate Suzuki, Woojin Kang, Tomoka Ema, Kento Morimoto, Kanako Kato, Tomoko Matsumoto, Nanami Masuyama, Yusuke Kijima, Toshifumi Morimura, Fumihiro Sugiyama, Satoru Takahashi, Eiji Mizutani, Knut Woltjen, Nozomu Yachie, Masatsugu Ema

**Author notes:** Correspondence should be addressed to: Seiya Mizuno, Laboratory Animal Resource Center, Trans-border Medical Research Center, University of Tsukuba, Tsukuba, Ibaraki 305-8575, Japan Masatsugu Ema, Shiga University of Medical Science, Seta, Tsukinowa-cho, Otsu, Shiga 520-2192, Japan. These authors contributed equally to this work (Eiichi Okamura, Seiya Mizuno, Shoma Matsumoto).

## Abstract

Pronuclear microinjection is the most popular method for producing transgenic (Tg) animals. Because the production efficiency is typically less than 20%, phenotypic characterization of Tg animals is generally performed on the next generation (F_1_) onwards. However, apart from in rodents, in many animal species with long generation times, it is desirable to perform phenotyping in the founder (F0) generation. In this study, we attempted to optimize a method of Tg mouse production to achieve higher Tg production efficiency using piggyBac transposon systems and established optimal conditions under which almost all individuals in the F0 generation were Tg. We also succeeded in generating bacterial artificial chromosome Tg mice with efficiency of approximately 70%. By combining this method with genome editing technology, we established a new strategy to perform phenotyping of mice with tissue-specific knockout using the F0 generation. Taking the obtained findings together, by using this method, experimental research using Tg animals can be carried out more efficiently.

## Introduction

Transgenic (Tg) technology enables genetic engineering techniques to introduce foreign DNA sequences, namely transgenes from other species or artificial ones, into the genomes of organisms. This technology can be used to produce organisms that express useful traits, such as resistance to infectious diseases, a high growth rate, and herbicide resistance. In the medical field, Tg technology has also made significant contributions, including the creation of disease model animals and the elucidation of gene function^1–4^. Another method for inserting foreign DNA into the genome is the knock-in (KI) approach, which involves inserting foreign genes into specific targeted locations, in contrast to Tg technology in which genes are inserted into the genome at random. Although KI technology has the advantage of being able to perform genetic modification in a predictable manner, it is generally inefficient and has difficulty introducing large-sized DNA. These problems with KI technology are being overcome in some animals such as mice and rats via the development of genome editing technology^5–7^. However, Tg technology is still considered useful for introducing foreign genes into large animal species, including rabbit, pig, cattle, and non-human primates, because efficient KI technology for such purposes has not yet been established.

The first Tg animal reported globally was a Tg mouse created by Jaenisch and Minz in the 1970s^8^. A retrovirus was used as a vector for producing this mouse, but the transgene was not expressed persistently due to silencing. In 1980, a pronuclear microinjection method, which is currently the most popular technique for producing Tg mice, was reported^9^. However, the efficiency of generating Tg mice using this method is approximately 20% at best^10^. Therefore, obtaining enough Tg mice in the F0 generation is difficult, and thus phenotypic analysis is generally performed using animals from the next generation onwards. The efficiency of Tg animal production should thus be improved in order to enhance research efficiency and to reduce the number of animals used, from the perspective of animal welfare. A more efficient method for generating Tg mice is a method using lentivirus first reported in 2003^11^, in which Tg mice can be produced with approximately 80%–100% efficiency. However, the lentiviral method is associated with the limitation that it can introduce transgenes of approximately 8 kbp at most.

Another method for generating Tg animals is the transposon-based method. Transposons are DNA sequences that can move from one location to another within the genome of an organism, and were originally found in maize^12^. During the transposition, the transposase enzymes bind to the inverted terminal repeats (ITRs) that define the boundaries of the transposon and cut out the DNA sequence between ITRs from the genome and insert it into another location. Several transposon-based methods have been established and used to produce Tg animals, including Tol2^13^, Sleeping Beauty^14^, and the piggyBac^15^ system. Transposon-based methods can introduce large transgenes of 100 kbp or more, but the Tg production efficiency is generally less than 45%^16^, which is lower than for the lentiviral method.

Recent developments in genome editing technology have made it possible to very efficiently produce systemic knockout (KO) animals, in which large-scale phenotypic screening analysis at the F0 generation can be conducted^17^. Approximately 23% of gene KOs result in embryonic lethality^18^, in which case tissue-specific KOs are often required to explore gene functions in a given tissue. However, the establishment of tissue-specific Cre Tg mice and flox mouse strains is time-consuming and laborious. Although many such mouse strains have been created and useful bioresources have been established, tissue-specific KO often takes more than a year to obtain, cross, and finally obtain the next generation for phenotypic analysis. Therefore, several methods have been reported to perform tissue-specific KO in a single generation^19–25^, but each method has its own advantages and disadvantages, and thus they should be used appropriately depending on the purpose.

Here, we report a novel strategy for generating tissue-specific KO called ScKiP (single-step cKO mouse production with piggyBac). We increased the efficiency of generating Tg mice with the piggyBac transposon system by optimizing the method to deliver transposase and donor DNA into the mouse zygote, and succeeded in producing Tg mice with an efficiency of nearly 100%. This method even enabled us to introduce an approximately 170 kbp bacterial artificial chromosome (BAC) vector with an efficiency of approximately 70%. By using this more efficient method, we succeeded in establishing a new approach to performing tissue-specific KO at the F0 generation. This approach is expected to contribute to animal welfare by reducing the number of animals used and also improve the efficiency of research.

## Materials and methods

### Study approval

All animal experiments were carried out in accordance with the Fundamental Guidelines for Proper Conduct of Animal Experiments and Related Activities in Academic Research Institutions under the jurisdiction of the Ministry of Education, Culture, Sports, Science and Technology, and Guidelines for Proper Conduct of Animal Experiments from the Science Council of Japan. All animal experimental procedures were approved by the Animal Care and Use Committee of Shiga University of Medical Science (2020-6-21) and the Institutional Animal Experiment Committee of the University of Tsukuba (22-019).

### Mice

C57BL/6J and ICR mice were purchased from Charles River Laboratories (Yokohama, Japan). PWK/Phj mice (RBRC00213) were obtained from RIKEN BioResource Research center (Tsukuba, Japan). The mice were maintained in plastic cages under pathogen-free conditions in a room maintained at 23.5 ± 2.5°C and 52.5 ± 12.5% relative humidity under a 14 h light:10 h dark cycle.

### Establishment of CdEC flox mouse line

We selected a sequence (5′-CGC CCA TCT TCT AGA AAG AC-3′) located in the first intron of the *Gt(ROSA)26Sor* as the CRISPR target. Briefly, an expression cassette consisting of CAG promoter, floxed EGFP, and Cas9NmC was placed between the 5′-homology arm and the 3′-homology arm. A rabbit beta-globin polyadenylation signal sequence was placed downstream of each of floxed EGFP and Cas9NmC. Cas9NmC is a fusion protein of Cas9 and part of mouse Cdt1 that we previously reported^26^. The CRISPR–Cas9 ribonucleoprotein complex and donor DNA were microinjected into zygotes of C57BL/6J mice, in accordance with our previous report^27^. Subsequently, microinjected zygotes were transferred into the oviducts of pseudopregnant ICR female and newborns were obtained. Genotyping for confirmation of the KI allele and random integration allele was performed using the same methods as in our previous report^26^. The CdEC flox mouse line was deposited to RIKEN BioResource Research center (RBRC12114).

### *In vitro* fertilization, electroporation, and microinjection

Fertilized eggs were obtained by *in vitro* fertilization using a method described elsewhere^28^. Briefly, oocytes were collected from oviducts of 10–14-week-old C57BL/6J females superovulated by the intraperitoneal administration of CARD HyperOva (Kyudo, Tosu, Japan), followed by human chorionic gonadotropin (hCG). Sperm were collected from the caudal epididymis of 12–16-week-old C57BL/6J males, and then preincubated in Fertiup Mouse Sperm Preincubation Medium (Kyudo). Insemination was performed in CARD medium (Kyudo), followed by incubation at 37 °C in an atmosphere containing 5% CO2 for 3 to 6 h. Fertilized eggs were washed to remove cumulus cells and sperm, and incubated in KSOM medium (Ark Resource, Kumamoto, Japan) until electroporation. Electroporation of fertilized eggs was performed in Opti-MEM I medium containing piggyBac mRNA using Super Electroporator NEPA21 (Nepa Gene, Ichikawa, Japan) as described elsewhere^29^ with minor modifications. In this study, the poring pulse was set as follows: 225V, 2 ms pulse width, 50ms pulse interval, 4 pulses, 10% attenuation rate, and + polarity. Meanwhile, the transfer pulse was set as follows: 20V, 50ms pulse width, 50ms pulse interval, 5 pulses, 40% attenuation rate, and ± polarity. The eggs were incubated until pronuclei were clearly visible. The transposon DNA was microinjected under a microscope equipped with a micromanipulator and Femtojet (Eppendorf, Hamburg, Germany). The injected one-cell embryos were cultured in KSOM medium until the two-cell stage and then transferred into pseudopregnant ICR mice.

### *In vitro* transcription

The template plasmid for piggyBac transposase (mPBase) for *in vitro* transcription was made by replacement of a mouse codon-optimized mPBase cDNA sequence with an EGFP sequence of pcDNA3.1-EGFP-poly(A)^30^. After linearization of this template plasmid using XhoI restriction enzyme, mPBase mRNA was synthesized using the mMESSAGE mMACHINE T7 ULTRA Transcription Kit (AM1345; Thermo Fisher Scientific, Waltham, MA, USA) or mMESSAGE mMACHINE T7 Transcription Kit (AM1344; Thermo Fisher Scientific).

### BAC DNA recombination

A BAC DNA clone (RP24-125B24) containing the mouse *Flk1* locus was obtained from Invitrogen (Invitrogen, Carlsbad, CA, USA). Modification of the BAC DNA to insert the GFP sequence into the first exon of the *Flk1* gene was performed in a previous study^31,32^. To insert ITR (inverted terminal repeat) sequences and Kusabira-Orange expression cassette into the backbone vector region, the BAC DNA was further modified by the same method as described in that previous study^31^ using the RED/ET recombination technique (Gene Bridges, Heidelberg, Germany). Collected recombinants were identified by screening for kanamycin resistance, followed by PCR analysis. The PGK-gb2-neo expression cassette was not excised.

### Genotyping of the embryo

E11.5 (embryonic day 11.5) stage embryos were collected from the uteri of surrogate mothers after euthanasia and the placenta and yolk sac were removed. The fluorescent signal was captured under a fluorescent stereoscopic microscope (FL5; Leica Microsystems, Wetzlar, Germany) equipped with a camera (DP73; Olympus, Shinjyuku, Japan). Genomic DNA from the embryos was extracted by phenol/chloroform extraction. Genotyping PCR was performed using a primer set to detect the GFP sequence using conventional PCR (T100; Bio-Rad, Hercules, CA, USA). To determine the copy number of the transgene, droplet digital PCR (ddPCR) was performed using the Qx200 Droplet Digital PCR system (Bio-Rad) with the GFP and Tfrc Primer/Probe set.

### Southern blotting

Genome DNA was extracted from mouse embryos with phenol and chloroform, and then purified by ethanol precipitation. After the digestion of 5µg of genomic DNA by EcoRI, the concentration was measured using a Quantus Fluorometer (Promega, Madison, WI, USA). One microgram of digested DNA was electrophoresed in 1% agarose gel. The DNA was blotted onto a nylon membrane (Amersham Hybond-N^+^; GE Healthcare, Chicago, IL, USA) and hybridized with DIG-labeled probe using DIG Easy Hyb (Roche Diagnostics, Mannheim, Germany). Salmon sperm DNA was used in the blocking step. After the reaction with anti-digoxigenin-AP Fab fragments (Roche), the bands were detected by enhanced chemiluminescence using CDP-Star (Roche) and ImageQuant LAS 4000 Mini (GE Healthcare). All probes were synthesized by PCR using DIG DNA Labeling Mix, 10× Conc. (Roche).

### Inverse PCR

PCR was performed using a method described elsewhere^33^ with minor modifications. In this study, 3µg of DNA was digested by EcoRI and the digested DNA was self-ligated using DNA Ligation Kit (Mighty Mix; Takara Bio, Kusatsu, Japan) after purification by ethanol precipitation. Regions where transgenes became integrated were amplified using transgene-specific primers by shuttle PCR. Nested PCR was also performed to enrich the genomic regions of interest.

### Echocardiography

A Vevo 2100 High-Resolution Imaging System (Visual Sonics Inc, Toronto, ON, Canada) was used to evaluate cardiac contractility. Mice were anesthetized with 2.5% isoflurane in oxygen and placed on a warmed platform. The isoflurane concentration was maintained at 1.5% during imaging. An ultrasound gel was applied to the left anterior thorax after the removal of fur from the body. Short-axis M-mode images were obtained at the level of the papillary muscle using a 40-MHz transducer. Left ventricular end-diastolic diameter (LVEDD) and left ventricular end-systolic diameter (LVESD) were measured using Vevo LAB software (FUJIFILM Visualsonics, Toronto, ON, Canada). Fractional shortening (FS) was calculated at three different time points of each mouse as follows: FS (%) = (LVEDD – LVESD)/LVEDD × 100.

## Results

### piggyBac transgenesis by electroporation and pronuclear microinjection

Although previous works achieved considerably high efficiency in the generation of Tg mice^15^, we assumed that Tg mice could be generated with even higher efficiency by initiating the expression of mammalian codon-optimized mPBase at an earlier timing than in the previous methods. To test this, we introduced *in vitro-*synthesized mPBase mRNA into zygotes by electroporation early after *in vitro* fertilization (Figure 1a). Consistent with previous reports describing that excessive amounts of mPBase inhibit transposition activity, a phenomenon called overproduction inhibition (OPI), we confirmed that OPI occurred at a high dose of mPBase in mouse ES cells, by varying the amount of mPBase expression plasmid with a certain amount of donor plasmid DNA (Figure 1b). To optimize the concentration of mPBase mRNA for electroporation, we applied it at 0, 20, 100, and 500ng/μl. After electroporation, the fertilized eggs were cultured *in vitro*, and donor plasmid DNA containing a GFP expression cassette (Figure 1c) was microinjected into the pronucleus at the PN3–4 stage (Figure 1a). Those embryos were cultured *in vitro* until the E4 stage. We counted the blastocysts exhibiting GFP fluorescence under each set of conditions by fluorescence microscopy and found that conditions with 500ng/μl mRNA gave the best results, at which approximately 80% of the blastocysts exhibited fluorescence (Figure 1d, e). To confirm this result at a later developmental stage, we transplanted two-cell-stage embryos into the oviduct of a surrogate mother, removed embryos from the uterus at E11.5 and dissected them, and conducted fluorescence observation as well as ddPCR to detect the transgene. The obtained results indicated that, for the 100ng/μl group, 9 out of 10 were Tg, while for the 500 and 1000ng/μl groups, all embryos were Tg. Although there were no significant differences in copy number of the transgene among all groups, the 500ng/μl group had the highest mean copy number (Figure 1f). From these results, we concluded that an mRNA concentration of 500ng/μl is optimal to produce Tg mice.

**Figure 1.**
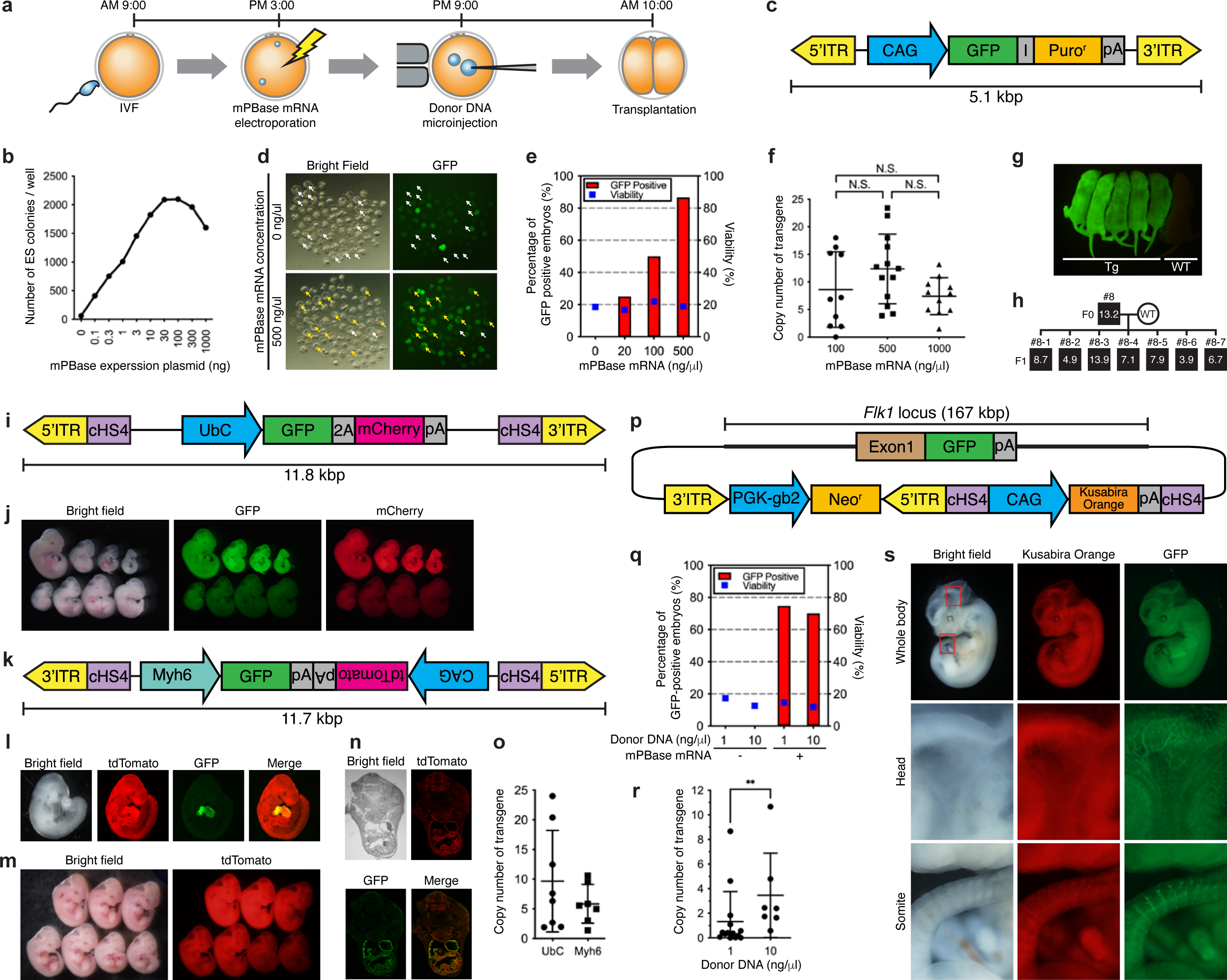
Generation of Tg mice by mPBase mRNA electroporation and donor plasmid DNA pronuclear microinjection (a) Schematic representation of the method to introduce mPBase mRNA and donor plasmid DNA. Typical schedule for performing each procedure is shown at the top of the figure. (b) Donor plasmid introduction into the ES cell genome using the piggyBac transposon system. ES cells were transfected with different concentrations of the mPBase expression plasmid vector and the same concentration of the donor plasmid, and the number of puromycin-resistant colonies was counted. (c) Schematic representation of CAG-EGFP-IRES-Puro donor plasmid DNA. Only the region between two ITRs is described. (d, e) Investigation of the optimal concentration of mPBase mRNA by observing blastocyst embryos using fluorescence microscopy. Fluorescence microscopic images of conditions with mRNA at 0 and 500ng/μl are shown in d. Tg embryos were judged by the presence of GFP fluorescence (orange arrow: GFP positive, white arrow: GFP negative blastocysts), with GFP-positive rates being displayed as a bar graph in e. Blue dots indicate blastocyst development rates for the number of fertilized eggs tested. (f) Investigation of the optimal concentration of mPBase mRNA using DNAs from E11.5 embryos. Transgene copy numbers were determined by ddPCR for the GFP sequence. (g) Fluorescent image of born Tg mice at the P1 (postnatal day 1) stage. (h) A mating test was performed to confirm germline transmission of the transgene. Three Tg male mice (#8, #9, #10) were crossed with wild-type (WT) females; the pedigree of line #8 is shown. Regarding line #8, copy numbers of the transgene were determined by ddPCR using DNA obtained from the tail tip. The numbers in black boxes indicate the copy numbers of the transgene determined by ddPCR using tail-derived DNA from F0 and F1 animals. (i) Schematic representation of UbC pro.-GFP-P2A-mCherry donor plasmid DNA. Only the region between two ITRs is described. cHS4: chicken hypersensitive site-4 insulator element. (j) Fluorescent images of E11.5 embryos harboring UbC pro.-GFP-P2A-mCherry transgene. Bright field (left), GFP fluorescence (middle), and mCherry fluorescence images (right) are shown. (k) Schematic representation of Myh6 pro.-GFP-CAG-tdTomato donor plasmid DNA. Only the region between two ITRs is described. (l) Fluorescent images of representative E9.5 embryos harboring Myh6 pro.-GFP-CAG-tdTomato transgene. Bright field image taken using a stereomicroscope (left), tdTomato fluorescence (2^nd^ from left), GFP fluorescence (2^nd^ from right), and merged images (right) taken using a confocal microscope are shown. (m) Fluorescent images of E11.5 embryos harboring Myh6 pro.-GFP-CAG-tdTomato transgene. Bright field (left) and tdTomato fluorescence images (right) are shown. (n) Confocal microscopic images of representative E11.5 embryos harboring Myh6 pro.-GFP-CAG-tdTomato transgene. Transverse frozen sections were prepared. Bright field image taken using a phase contrast microscope (upper left), tdTomato fluorescence (upper right), GFP fluorescence (lower left), and merged images (lower right) taken using a confocal microscope are shown. (o) Copy number analysis of Tg embryos harboring UbC pro.-GFP-P2A-mCherry transgene (UbC) and Myh6 pro.-GFP-CAG-tdTomato transgene (Myh6). (p) Schematic representation of BAC DNA used as a donor vector. (q) Investigation of the optimal concentration of donor DNA using DNAs from E11.5 embryos. Transgenes were detected by conventional PCR and GFP-positive rates are displayed as a bar graph. Blue dots indicate E11.5 development rates for the number of transplanted two-cell-stage embryos. (r) Copy number analysis of BAC Tg. Transgene copy numbers were determined by ddPCR for the GFP sequence using DNAs from E11.5 embryos. (s) Fluorescent images of a representative E11.5 BAC Tg embryo. Bright field (left), Kusabira-Orange fluorescence (middle), and GFP fluorescence images (right) of the whole body (top), head (middle), and somite (bottom) region are shown. The red rectangular frames inside the whole body represent the head and somite region.

To confirm that this method would result in healthy births, we transplanted the fertilized eggs into surrogate mothers. We found that, out of 70 fertilized eggs, 12 (17%) animals were born; of these, 1 animal was eaten by its mother, while the rest of them (11 animals) were Tg (Figure 1g). To evaluate the reproductive ability of the F0 generation, three males were mated and all resulting offspring were found to be Tg (Figure 1h).

### Bacterial artificial chromosome DNA integration by piggyBac transposase

Relatively short plasmids were used as the donor DNA in our assay, and therefore we attempted to use larger donor DNAs with several expression cassettes. We constructed two plasmids with a larger 12kbp transgene. One was a construct containing an expression cassette for GFP-P2A-mCherry linked to the Ubiquitin C promoter (Figure 1i). All eight E11.5 embryos showed GFP and mCherry expression (Figure 1j). The other was a construct containing expression cassettes for GFP linked to the *Myh6* promoter and tdTomato linked to the CAG promoter (Figure 1k). We obtained four embryos that reached the E9.5 stage, seven that reached E11.5, and three newborns, all of which showed tdTomato expression (Figure 1l&m). We also confirmed heart-specific GFP expression in all of the embryos at E9.5 and E11.5 by fluorescence microscopy (Figure 1l&n). Finally, ddPCR was performed using DNAs extracted from E11.5 embryos to analyze the transgene copy number. The results revealed 9.7±8.6, and 5.8±3.3 copies of the transgenes in UbC pro.-GFP-P2A-mCherry and Myh6 pro.-GFP-CAG-tdTomato mice, respectively (Figure 1o).

Mice of 12kbp Tg were established at 100% efficiency, and therefore we applied this method to establish BAC Tg mice because the efficiency of BAC Tg mouse production is typically very low. In a previous study, we constructed BAC DNA with a GFP sequence linked to the *Flk1* promoter. Here, the ITR sequences and Kusabira-Orange cassette were inserted into the vector backbone of the BAC DNA (Figure 1p). The engineered BAC DNA was used as a donor DNA at two different concentrations (1 and 10ng/μl). Owing to the high viscosity of BAC DNA, microinjection at concentrations higher than 10ng/μl was not feasible. We obtained 14 Tg animals out of 19 embryos at E11.5 under conditions with 1ng/μl, while 7 out of 10 embryos were Tg at E11.5 under conditions with 10ng/μl only when mPBase had been introduced (Figure 1q). Thus, there was no significant difference in the percentage of Tg mice under the two sets of conditions. Next, we evaluated the transgene copy number in these Tg mice by ddPCR and found that the mean copy numbers were 0.39 and 2.39 for 1 and 10ng/μl, respectively, which were significantly different (Figure 1r). The BAC Tg embryos exhibited ubiquitous Kusabira-Orange expression throughout the body and specific GFP signals in the blood vessels (Figure 1s), consistent with a previous report^31^. Finally, PCR genotyping analysis indicated that the percentage of Tg mice among the offspring was very high (52 out of 94; 55%).

### Donor DNA injection into fertilized eggs refrigerated after electroporation

Although the method using electroporation and microinjection developed in this study can produce Tg mice with very high efficiency, the experimental schedule is inconvenient due to microinjection being performed late at night. Therefore, we modified the schedule by storing the eggs after electroporation in a fridge, and then performed microinjection the next morning (Figure 2a). After either a mixture of BFP/EGFP/tdTomato expression vectors (three colors) or a mixture of BFP/EGFP/tdTomato/E2Crimson (four colors) (Figure 2b) was microinjected, the fertilized eggs were transplanted into the oviduct of surrogate mothers at the one-cell stage. We dissected the embryos at E14.5 and checked the presence of fluorescence as an indicator of Tg. We obtained 12 embryos, 11 (91.7%) of which were Tg under the three-color conditions, while 3 out of 4 embryos (75%) were Tg under the four-color conditions (i.e., at least one fluorescence signal was shown) (Figure 2c&d). These results indicate that Tg mice can be generated at high efficiency even after refrigeration of the fertilized eggs.

**Figure 2.**
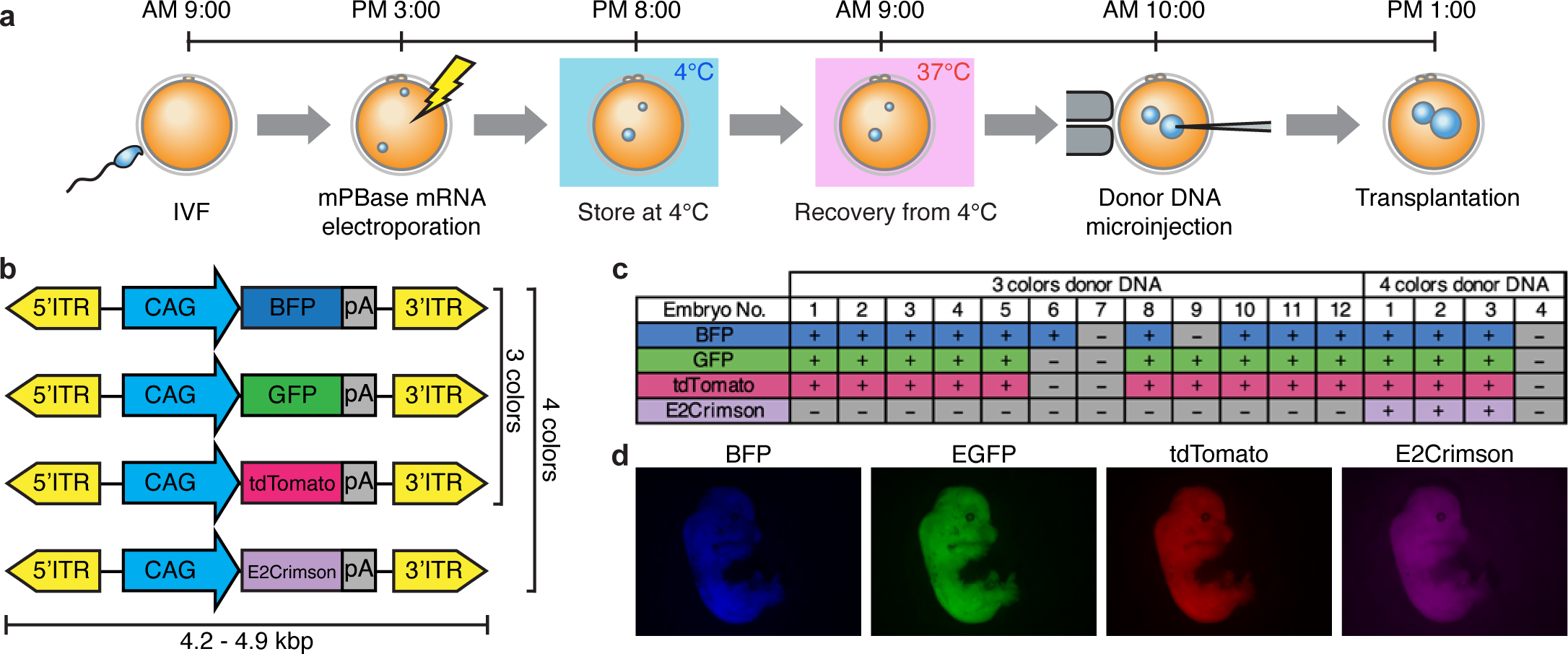
Generation of Tg mice using fertilized eggs refrigerated after electroporation (a) Schematic representation of method to introduce mPBase mRNA and donor plasmid DNA. Fertilized eggs were temporarily refrigerated after electroporation. Typical schedule for performing each procedure is shown at the top of the figure. (b) Schematic representation of BFP, EGFP, tdTomato, and E2Crimson expression cassettes on the donor plasmids. (c) Summary of the results of observing the obtained embryos by fluorescence microscopy. The presence of each fluorescence color is indicated by + and its absence is indicated by −. The results of three- (top) and four-color donor plasmid injection (bottom) are shown separately. (d) Fluorescent images of representative E11.5 embryo harboring four-color transgenes. BFP (left), EGFP (2^nd^ from left), tdTomato (2^nd^ from right), and E2Crimson fluorescent images (right) are shown.

### Rapid phenotyping of conditional KO mice at the F0 generation by ScKiP

Currently, for phenotypic analysis of essential genes in a certain tissue that play roles in embryonic survival, the Cre-loxP system must be used, which is time-consuming and laborious. By using the method established in this study, we attempted to establish a new strategy to generate conditional KO at the F0 generation by using piggyBac transgenes. First, we targeted the *Prmt1* gene, which causes dilated cardiomyopathy of young mice upon heart-specific KO, while its systemic KO is embryonically lethal^34^. For heart-specific KO, we constructed a donor plasmid with the Cre gene linked under a cardiac-specific *Myh6* promoter and a gRNA for the *Prmt1* gene expression cassette (Figure 3a). Prior to the conditional KO experiment, to check that the Myh6 promoter causes Cre expression in the heart, we produced Tg mice in the GRR mouse^35^ strain background. The GRR allele has a structure in which the floxed EGFP-polyA sequence and the tdTomato gene are linked under the CAG promoter, and tdTomato is expressed following Cre-mediated excision (Figure 3b). The donor plasmid was microinjected into the pronuclei after storage at 4°C overnight, and those eggs were transplanted into the oviduct of a surrogate mother at the one-cell stage. We obtained 14 offspring, 10 of which were revealed to possess the transgene (GRR KI/Tg+) by genotyping PCR analysis. Fluorescent microscopy of the born mice indicated that most of the GRR KI/Tg+ mice showed strong tdsRed fluorescence in the heart, along with nonspecific expression in regions including the skin (Figure 3c&d). To perform conditional KO, we used the CdEC flox mouse strain containing the KI allele in which the floxed EGFP-polyA sequence and the Cas9NmC gene are linked under the CAG promoter, and Cas9NmC is expressed only after Cre expression (Figure 3e). Tg mice with the same donor vector were produced in a similar way in the CdEC flox mouse strain. The results revealed that 10 offspring were obtained, and genotyping PCR analysis revealed that all animals possessed the CdEC flox KI allele, while only one had the transgene (CdEC KI/Tg+). Echocardiography at 2, 4, and 6 weeks old revealed that cardiac contractility of the CdEC KI /Tg+ mouse was reduced compared with that in mice without transgene (CdEC KI/Tg−, Figure 3f). At 50 days old, we noticed that the CdEC KI/Tg+ mouse was sluggish and wheezed, and thus euthanized it. The heart of this CdEC KI/Tg+ mouse weighed 188mg, which was markedly heavier than that of a CdEC KI/Tg− mouse (101mg, Figure 3g), suggesting dilated cardiomyopathy similar to that of conventional cKO mice reported previously^34^. Thus, although we were able to observe a phenotype of heart-specific KO of the *Prmt1* gene at the F0 generation, the rate of Tg mice relative to all offspring was low at 10% in this experiment. This may have been due to ectopic Cre expression other than in the heart in Tg embryos, resulting in embryonic lethality.

**Figure 3.**
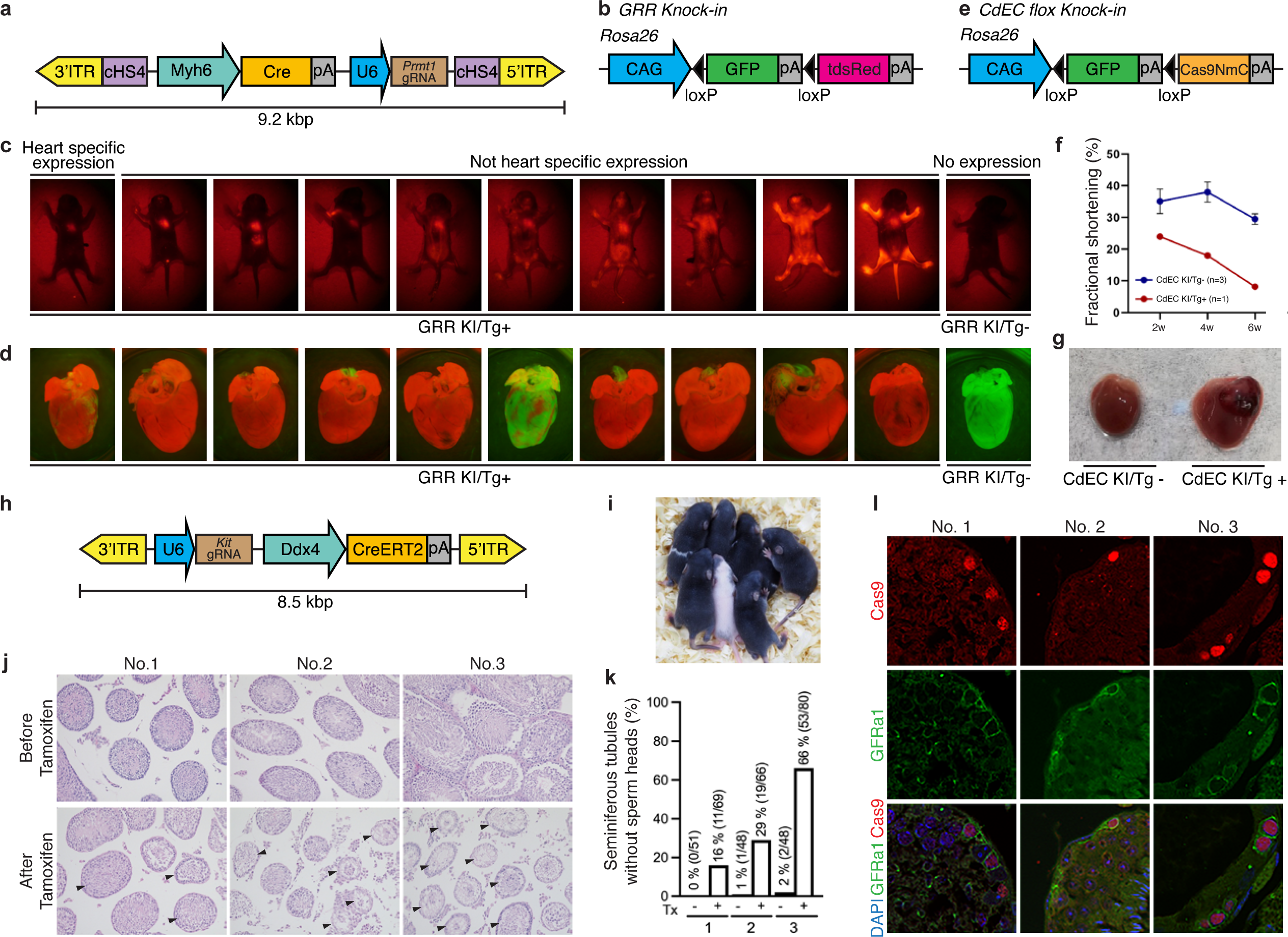
Rapid phenotyping of conditional KO mice (a) Schematic representation of Myh6-Cre-Prmt1 gRNA donor plasmid DNA. Only the region between two ITRs is described. cHS4: chicken hypersensitive site-4 insulator element. pA: polyadenylation signal. (b) Schematic representation of GRR KI allele. (c) Fluorescent images of born GRR KI mice with Myh6-Cre-Prmt1 gRNA transgene (GRR KI/Tg+) at the P4 (postnatal day 4) stage. Fluorescent image of GRR KI mice without the transgene (GRR KI/Tg-) was shown as a control. (d) Fluorescent images of heart from GRR KI/Tg+ and GRR KI/Tg- mice. (e) Schematic representation of CdEC flox KI allele. (f) Results of echocardiography on 2-, 4-, and 6-week-old CdEC flox KI mouse with Myh6-Cre-Prmt1 gRNA transgene (CdEC KI/Tg+) or without transgene (CdEC KI/Tg-). (g) Morphogenic observation of hearts from 6-week-old CdEC KI/Tg+ and CdEC KI/Tg- mice. (h) Schematic representation of gRNA-Ddx4-CreERT2 donor plasmid DNA. Only the region between two ITRs is described. (i) Observation of newborn CdEC KI mice with the gRNA-Ddx4-CreERT2 transgene (CdEC KI/Tg+). (j) Results of HE-staining analysis of testes removed from CdEC KI/Tg+ mice before and after tamoxifen injection. Arrowheads indicate seminiferous tubules not containing sperm heads. (k) Percentage of seminiferous tubules without sperm head in the testes from CdEC KI/Tg+ mice before and after tamoxifen injection. (l) Results of the immunostaining analysis of testes from CdEC KI/Tg+ mice after tamoxifen injection using anti-Cas9 and anti-GFRα1 antibody.

Given the success of characterizing the heart-specific phenotype associated with *Prmt1*, we applied this strategy to characterize mice with germ cell-specific KO of the *Kit* gene at the F0 generation. It has been reported that mice with hypo-functional *Kit* point mutant (*W^V^*/*W^V^*) had no sperm, while systemic KO with deletion of the transmembrane domain (*W*/*W*) resulted in perinatal- or late fetal-stage lethality. To avoid lethality due to non-specific Cre-loxP recombination, we used germ cell-specific *Ddx4* promoter-driven CreERT2, which works only in the presence of tamoxifen as a component of the donor plasmid with an expression cassette of gRNA for the *Kit* gene (Figure 3h). Similar to the procedure in the previous experiment, after mPBase introduction into fertilized eggs prepared using CdEC flox KI male and wild-type female mice, the donor plasmid was microinjected into the pronuclei, followed by transplantation into the oviduct of a surrogate mother at the one-cell stage. We obtained 14 offspring, including 11 males and 3 females (Figure 3i), all of which were revealed to have the transgene and KI allele (CdEC KI/Tg+) by genotyping PCR. Nine males were used for subsequent experiments. From each of these nine males, one testis was removed as a control (before tamoxifen sample) at 16 to 17 weeks old. Tamoxifen was injected three times at 18, 22, and 23 weeks old to induce Cre-loxP recombination. During this process, six mice died probably due to tamoxifen toxicity, postoperative complications, or non-specific *Kit* gene KO in cells other than germ cells. At 32 weeks old, the remaining three males were euthanized and their testes were collected (after tamoxifen sample). HE-stained sections from the removed testes were prepared and the proportion of seminiferous tubules not containing sperm heads was determined (Figure 3j). It was found that the number of seminiferous tubules without sperm increased after tamoxifen administration compared with that before tamoxifen administration in all three males (Figure 3k). When the percentage of Cas9-expressing undifferentiated spermatogonia after tamoxifen administration was examined by immunostaining using anti-Cas9 and anti-GFRα1 antibody, reductions in undifferentiated spermatogonia (8.5%, 86%, and 59%) were seen in samples No. 1, 2, and 3, respectively (Figure 3l). We named this cKO method “ScKiP,” which stands for single-step cKO mouse production with piggyBac.

### Preference of the Tg integration site and parental alleles in mouse zygote

To reveal the preference of integration sites into which Tg is introduced by our piggyBac system, we prepared fertilized eggs with PWK/Phj (PWK) strain sperm and C57BL/6J (B6) strain oocyte, and then injected mPBase mRNA and a donor plasmid DNA containing GFP and a DNA barcode sequence (Figure 4a). We obtained 17 E11.5 embryos, 9 of which were Tg as judged by fluorescence microscopy and genomic PCR. We investigated the number of integration sites in each Tg embryo by Southern blotting (Figure 4b). We confirmed that one to three transgenes had integrated into different genomic regions of the genome in each embryo. We further performed ddPCR analysis to evaluate the copy number (Figure 4c). Subsequently, we detected the DNA barcode sequence of the Tg by Amplicon-seq; overall, we detected 16 unique DNA barcode sequences. To investigate the sites of integration of the Tg in the genome, we performed inverse PCR and confirmed the results by genotyping PCR using primers specific for the DNA barcode sequence. Focusing on the genomic features of the integration site, 9 out of 16 transgene were located in introns, 3 were in repeat sequences, 3 were intergenic regions and only 1 was located coding sequence. (Figure 4d). Using information on the SNPs between the PWK and B6 genomes, we could determine whether the transgene was introduced into the paternal or maternal allele for 13 out of 16 transgenes (Figure 4e). Overall, 3 transgenes were on the maternal allele, even though we injected donor DNA into the male pronucleus. This suggested that the DNA integration event by mPBase continued after pronuclear fusion. To obtain a deeper understanding of the integration sites, we performed comparative analysis using ChIP-seq data. It was reported that the integration sites for mPBase were enriched in regions with H3K4me3^36^. We downloaded H3K4me3 ChIP-seq data of the zygote and early two-cell embryo with PWK sperm and B6 oocyte from the GEO database, and compared the peaks with the integration sites (GSE71434)^37^. Overall, 31.25% of integration sites overlapped with H3K4me3, which is generally located on the active TSS, for the integrated allele (Figure 4f, g).

**Figure 4.**
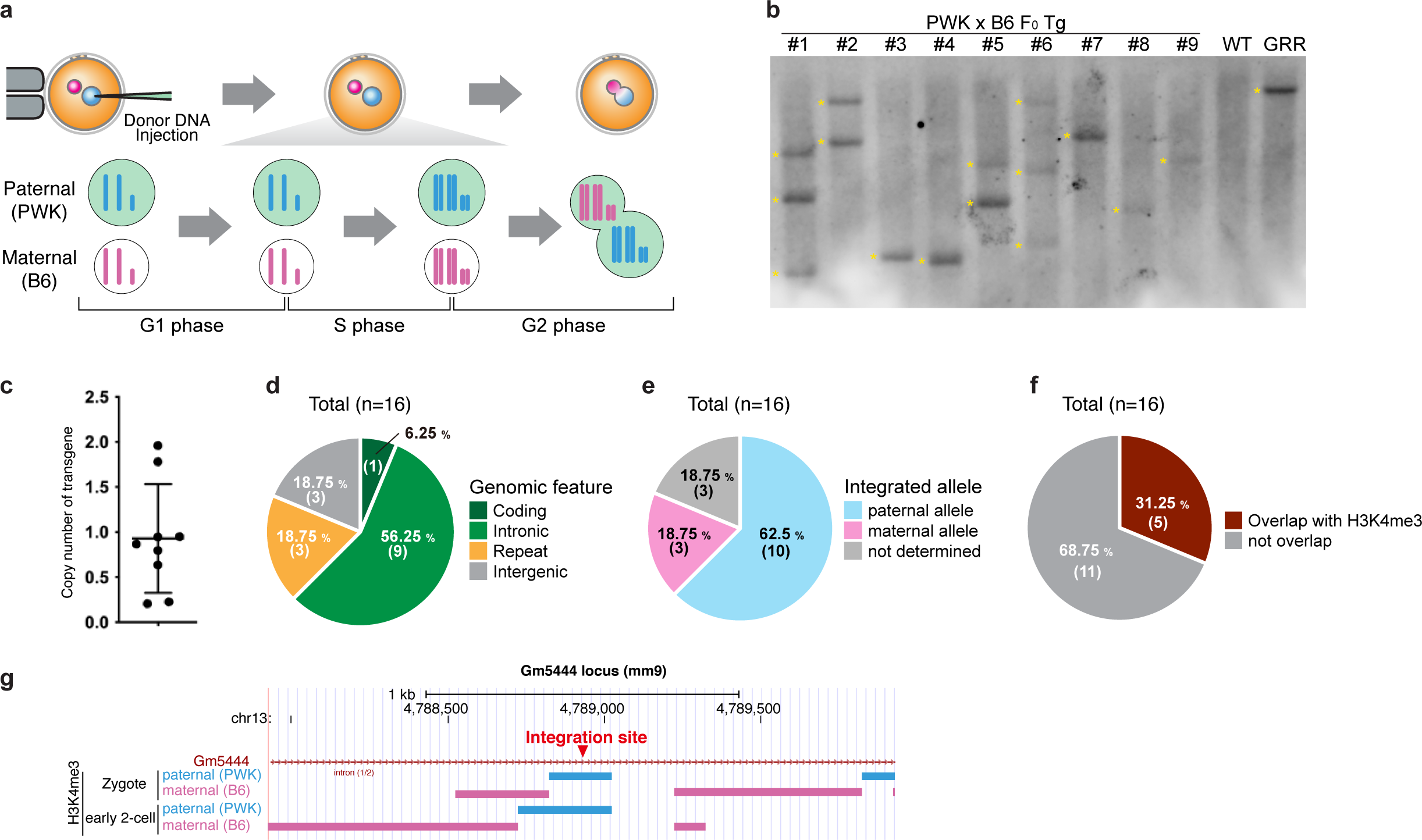
Generation of Tg mice using oocytes from C57BL6J and sperm from PWK strain (a) Schematic showing the timing of donor DNA microinjections relative to cell-cycle progression in the mouse embryo. (b) Results of Southern blot analysis using DNA from E11.5 Tg embryo obtained with a probe for the GFP region. DNA from WT mouse was used as a negative control. DNA from GRR KI mouse was used as a positive control. (c) Copy number analysis of the transgenes using DNA from E11.5 Tg embryo obtained by ddPCR with a probe for the GFP region. (d) Summary of genome-wide distribution of the transgene insertion sites. (e) Summary of the transgene integration alles. (f) Percentage of transgene insertion sites overlapping with H3K4me3 peaks. (g) An example of a transgene insertion region overlapping with H3K4me3 peaks.

## Discussion

In this study, we established a highly efficient Tg production method using the piggyBac transposon system. Three types of transposon systems are commonly used to create Tg animals: piggyBac, Sleeping Beauty, and Tol2^38^. Among them, we employed piggyBac because it is the only transposase used to create Tg mice with transgenes larger than 150 kbp^39^. By mPBase mRNA electroporation and pronuclear donor DNA microinjection, a plasmid-sized transgene with 100% efficiency and a BAC transgene with approximately 70% efficiency could be introduced.

Previous works indicated that the efficiency of Tg mouse production by mPBase was typically 20%–70%. The previously described methods and ours differ in terms of how mPBase is introduced. In several previous studies, mPBase expression plasmid vector was introduced into the cytoplasm of unfertilized eggs or the pronucleus of fertilized eggs^15,40–42^. Urschits *et al.* showed that mPBase protein expression was at the background level until 6 h after injection of the expression plasmid, and peak expression was observed at about 30 h^40^. Therefore, mPBase is fully expressed after the two-cell stage, and donor plasmids introduced at the one-cell stage might be degraded or diluted until the mPBase protein functions. In other reported studies, mPBase mRNA was used instead of an expression plasmid, but it was injected at the pronuclear stage^43–47^, and thus full expression may have occurred after the two-cell stage. In contrast, in this study, we attempted to introduce mPBase mRNA immediately after fertilization using electroporation. In this method, mPBase was sufficiently expressed when donor DNA was injected into the pronucleus and the transgene integration may have occurred efficiently before donor DNA degradation or dilution.

In this study, although we attempted to improve the efficiency of piggyBac transgenesis in mice, improvement of Tg production efficiency is also useful for other animal species. In rat, another popular experimental animal, pronuclear injection of a piggyBac expression plasmid vector and donor DNA has been used to generate Tg animals, but the efficiency was generally less than 30%^44,48,49^. Although a study reported that Tg rats can be produced more efficiently (33%–100%, mean of nearly 80%) using mPBase mRNA pronuclear microinjection^50^, it may be possible to consistently produce Tg animals even more efficiently if mPBase mRNA is introduced at an earlier timing after *in vitro* fertilization. Transgenic technology is also important for livestock, such as cattle, pig, and sheep, but there are very limited reports on application of the piggyBac transposon system to their zygotes^51,52^. Somatic cell nuclear transfer is one of the most common methods to create Tg livestock^53,54^. However, this approach is technically difficult, and the birth rate is typically low. Although no reports have yet been published on the use of mPBase mRNA injection to produce Tg livestock, this approach is worth pursuing.

Mice with tissue-specific conditional KO are very useful to elucidate gene functions *in vivo* but are time-consuming to produce because they typically require flox and Cre-Tg animal strains to be obtained and crossed. Recently, new methods have been developed to perform tissue-specific KO in a rapid manner. One example involves introducing CRISPR-Cas9 ribonucleoproteins into mice using nanoparticles, which enables efficient KO in several tissues, including liver, muscle, brain, kidney, and lung^19–22^. Another example involves introducing the flox allele into zygotes obtained from tissue-specific Cre-Tg mice by KI technology^23^. A third example involves the introduction of gRNA by an adeno-associated virus (AAV) vector into tissue-specific Cas9-expression Tg mice. Finally, a method has been developed involving the introduction of gRNA and a tissue-specific Cre expression vector by AAV into mice harboring a Cre-loxP reaction-dependent Cas9 expression cassette. Although these methods are very useful, they vary in terms of their targetable tissues, efficiency of cKO generation, and procedural difficulty. Thus, it is important to carefully examine these options and choose an appropriate method considering the purpose of the planned research. In this study, as another option to perform cKO experiments within a short time, we developed the ScKiP method and tested its usefulness. For rigorous evaluation, we selected *Prmt1* and *Kit* as target genes, for which a cKO experiment is necessary to reveal their functions after birth because their systemic KO is embryonically lethal. Although we confirmed that this method is not strictly tissue-specific, probably due to positional effects, we could observe the expected phenotype. Therefore, the ScKiP method is considered to be useful for rapid screening analysis of gene function. In addition, using the ScKiP method, KO cells can be identified using the disappearance of EGFP fluorescence as an indicator, which is a feature not found in other methods described above.

Taking the obtained findings together, by using the Tg production method that we have established, we can obtain a sufficient number of animals for phenotypic analysis in the F0 generation. Our methods are expected to contribute to animal welfare by reducing the number of animals used and enabling research to be carried out more efficiently.

## Acknowledgments

pcDNA3.1-EGFP-poly(A) plasmid was kindly provided by Dr. Kazuo Yamagata (KINDAI University). This work was supported by JSPS KAKENHI Grant Numbers 21K05988, 23H03860 (to E.O.), 20K22611 (to S.M.), and 21H02388 (to M.E.). We thank Central Research Laboratory at Shiga University of Medical Science and Single-cell Genome Information Analysis Core (SignAC) at WPI-ASHBi, Kyoto University, for their support. We also thank Edanz (https://jp.edanz.com/ac) for editing a draft of this manuscript. Finally, we thank all members of the Department of Stem Cells and Human Disease Models, Research Center for Animal Life Science, Shiga University of Medical Science.

## Notes

### Competing Interest Statement

The authors have declared no competing interest.

